# Self-consistent automatic retrieval of single cell rotation enables highly reliable holo-tomographic flow cytometry

**DOI:** 10.64898/2026.02.24.707756

**Authors:** Daniele Pirone, Lisa Miccio, Vittorio Bianco, Pietro Ferraro, Pasquale Memmolo

## Abstract

Holo-Tomographic Flow Cytometry (HTFC) is a powerful, label-free imaging technique that allows obtaining three-dimensional (3D) refractive index (RI) tomograms of individual flowing and rotating cells. Determining the unknown orientation of moving cells is crucial for tomographic reconstruction. There is significant potential to enhance previous methods by increasing accuracy, achieving full automation, and reducing the dependence on a priori information. Here, we present a novel self-consistent method for the accurate and automatic retrieval of unknown rotations of single cells. The proposed method retrieves the sequence of rotation angles by an iterative reprojection-based optimization algorithm of the reconstructed 3D RI tomogram. The experimental results demonstrate that the method proposed in this work achieves full automation and superior accuracy, thus surpassing the existing techniques. This advancement markedly improves the scalability of HTFC, enabling automatic and highly reliable analysis of single cells.

---

Label-free microscopy is revolutionizing biomedical optical imaging thanks to avoiding conventional exogenous labelling agents [1]. Among the others, Holographic Tomography (HT) is increasingly emerging as a three-dimensional (3D) Quantitative Phase Imaging (QPI) technique able to provide the volumetric distribution of the refractive index (RI) values of a biological specimen [2-9]. Recently we demonstrated Holo-Tomographic Flow Cytometry (HTFC), which combines the advantages of the object rotation HT scheme and imaging flow cytometry [10-13]. HTFC records, along a fixed beam direction, multiple digital holograms of the same cell while flowing and rotating along a microfluidic channel. In this way, the 3D RI tomograms of single cells in suspension can be reconstructed with quasi-isotropic resolution, unlike widespread illumination scanning HT configurations [4-6]. Moreover, unlike conventional object rotation HT configurations [7,8], HTFC potentially adds the high-throughput property typical of flow cytometry systems, which is crucial to carry out advanced single-cell analyses in 3D, label-free, and quantitative way. However, the promising potential of HTFC has not yet been fully exploited due to the need for retrieving the unknown cells’ rolling angles, which are required to reconstruct their 3D RI tomograms in addition to the corresponding quantitative phase maps (QPMs). The numerical methods originally developed at this aim required prior information about the object’s shape/RI [10] or exploited theoretical microfluidic models [14]. Convolutional neural networks have recently been trained to estimate the orientation of red blood cells [15], but they struggle to generalize to biological samples never seen during training. Currently, the matching-based method is the most employed one in HTFC, which has allowed reaching most of the HTFC results by reducing the dependence on a priori information [16]. However, there is still a need for increasing accuracy and achieving full automation. Here we propose a novel self-consistent reprojection-based method for retrieving the unknown rolling angles in HTFC. The proposed algorithm reprojects iteratively the 3D RI tomogram to find the best rolling angles sequence optimizing the tomographic reconstruction. Our reprojection-based method overcomes limitations of existing techniques, achieving superior accuracy and full automation, which consequently enables a substantial speedup in tomographic reconstruction for flow cytometry.

Let *xyz* be a Cartesian reference system within the microfluidic channel, where *z* is the optical axis. The microfluidic system setting can ensure that cells flow along the *y*-axis and rotate around the sole *x*-axis in a continuous and quasi-uniform way [16]. If the recording frame rate is set to perceive and quantify the cell rotation between subsequent frames, it can be safely supposed that the angular variation (around the *x*-axis) between two successive frames is proportional to the position variation (along the *y*-axis).

So far, in the matching-based method [16], the unknown rolling angles have been retrieved by exploiting the latter property as

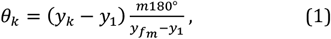

where *y*_*k*_ is the *y*-position at the frame *k* = 1, …, *K, m* is an integer number, and *f*_*m*_ is the frame at which a *m*180° rotation has occurred with respect to the first one *k* = 1 (*θ*_1_ is arbitrarily fixed at 0°). To implement this formula, the *f*_*m*_ frame must be identified. At this aim, the minimization of an ad hoc similarity metric needs to be performed, namely the Tamura Similarity Index (TSI), calculated between all the QPMs Ψ_*k*_ with *k* = 2, …, *K* and the first one Ψ_1_ [16]. For this reason, this method is referred to as matching-based rolling angles recovery method. For example, if *m* = 2, a full rotation (i.e., 360°) is searched. However, finding the correct *f*_*m*_ value often requires human verification for two main reasons. First, due to high noise and/or a quasi-spherical shape and/or a quasi-homogeneous inner content, the true *f*_*m*_ value may correspond to a local minimum near the global one. Second, multiple cell rotations can create several local minima, each tied to a different orientation. Moreover, even if the correct *f*_*m*_ frame is identified, angles are still corrupted by an intrinsic approximation error. In fact, Eq. (1) is assuming that the *m*180° rotation has been exactly sampled, but this is not true in most cases. In fact, usually a *m*180° + *ε* rotation is recorded at the *f*_*m*_ frame, instead of the exact *m*180° rotation. The *ε* error belongs to the interval 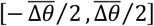, where 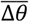 is the average angular sampling acquired by the HTFC system. Hence, the *ε* error propagates throughout the overall sequence of angles *θ*_*k*_ computed through Eq. (1). Instead, in the self-consistent reprojection-based rolling angles recovery method herein proposed, we formalize the proportionality between the angular variation (around the *x*-axis) and the translational variation (along the *y*-axis) in a different way, i.e. revisiting the Eq. (1) as

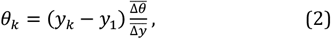

where 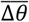 and 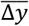 are the average angular increment and the average translational increment that occur between two successive frames. The 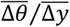 value is the average angular increment performed by the cell during its rotation for each pixel of translation along the *y*-axis. Hence, the formula in Eq. (2) is a generalization of the formula in Eq. (1), and they coincide if a *m*180° rotation is exactly acquired at the *f*_*m*_ frame.

We have employed a 3D numerical cell phantom (see Fig. 1) to assess our self-consistent reprojection-based rolling angle recovery method before application to real experimental data. To simulate an angular sequence, without loss of generality, we have fixed 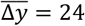 pixels. Then, starting from *y*_1_ = 0 pixels, we have created an array of *K* equidistant positions *y*_*k*_. To consider a non-uniform roto-translation, we have perturbed each position *y*_*k*_ by a *δ*_*k*_ value randomly drawn from the integer uniform distribution *U*(−6,6), with *k* = 2, …, *K*. Hence, we have computed *K* = 61 rolling angles *θ*_*k*_ through Eq. (2) by fixing 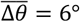 (average cell roto-translation of 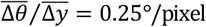). Note that the 360° rotation has not been sampled, as the cell passes from *θ*_60_ = 356° to *θ*_61_ = 361.5°. Finally, we have simulated the *K* QPMs Ψ_*k*_ by integrating the tomogram along the simulated *θ*_*k*_ angles, according to the linear approximation of the light propagation [11,16].

**Fig. 1.**
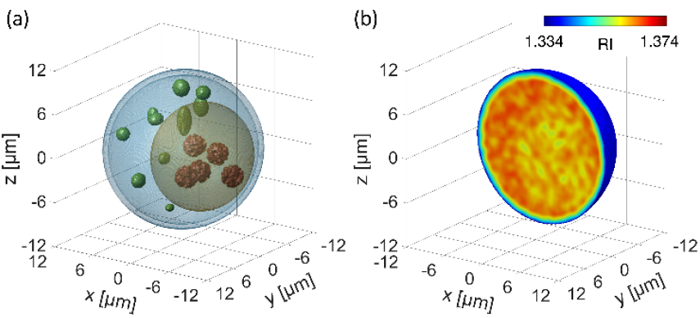
3D numerical cell phantom. **(a)** Isolevels representation of the cell, containing cell membrane (light blue), cytoplasm (blue), nucleus (yellow), 5 nucleoli (red), and 10 lipid droplets (green). **(b)** Central slice of the 3D RI tomogram.

Actually, in a real experimental problem,

1. the *K* QPMs Ψ_*k*_ are numerically extracted from the corresponding holographic region of interests (ROIs);
2. the *K* positions *y*_*k*_ are computed through the holographic tracking algorithms;
3. the average translational increment 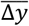 value is measured from the vector of positions *y*_*k*_ as

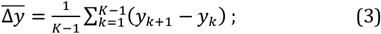
4. the average angular increment 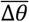 is estimated through the reprojection-based method;
5. the *K* unknown rolling angles *θ*_*k*_ are retrieved through Eq. (2).

Hereafter we discuss the reprojection-based method to estimate the 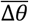 value at step 4 by using the *K* QPMs Ψ_*k*_, as also sketched in Fig. 2 throughthe 3D numerical cell phantom. It is based on the property that the tomogram *T*_*k*=1,…,*K*_, reconstructed by using all the QPMs, is rotated around the *x* -axis by an angle −Δ*θ*_1_ = −(*θ*_2_ − *θ*_1_) = −*θ*_2_ with respect to the truncated tomogram *T*_*k*=2,…,*K*_, reconstructed by using all the QPMs except the first one.

**Fig. 2.**
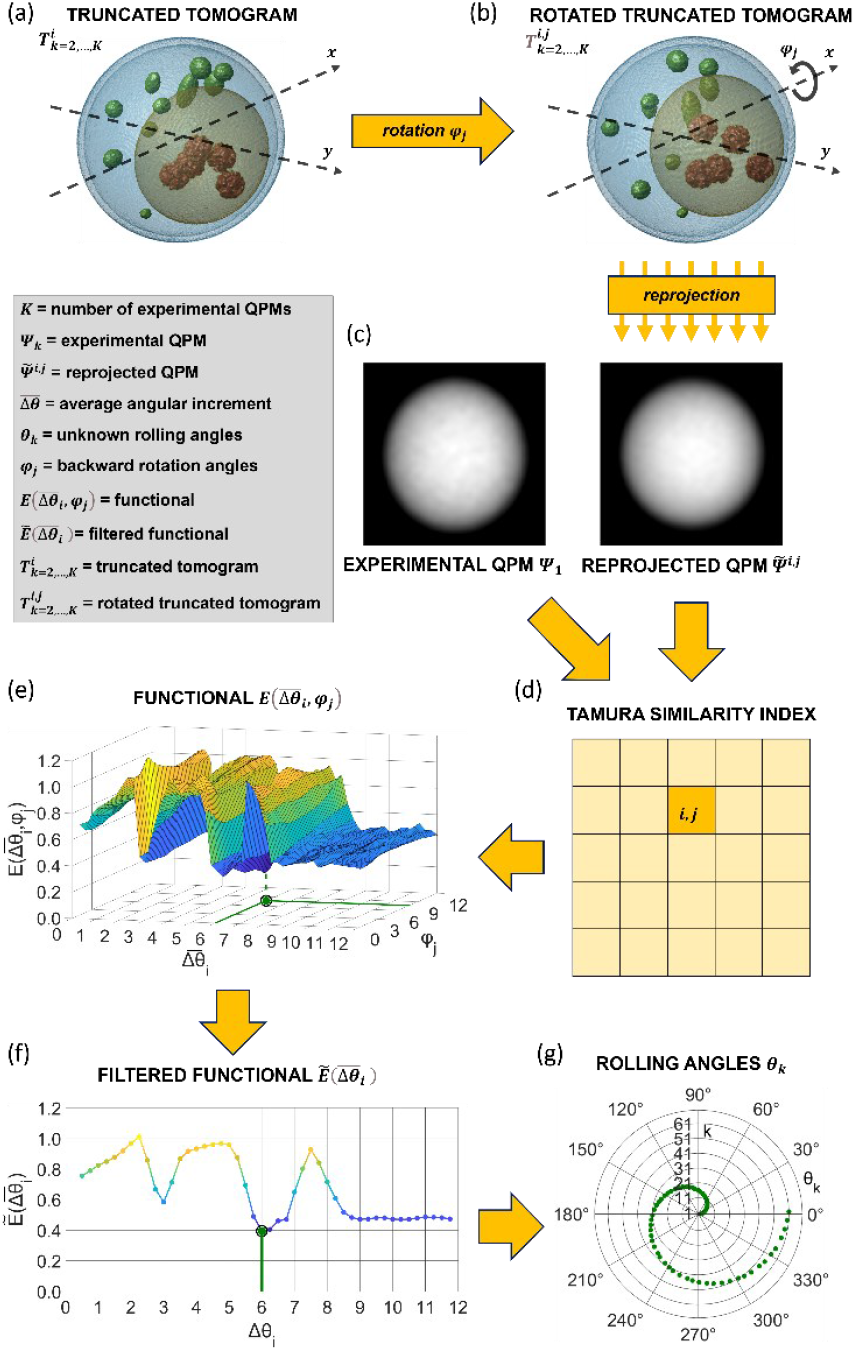
Workflow of the reprojection-based method. **(a)** The truncated tomogram 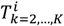, reconstructed at *N* average angular increments 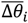, **(b)** is rotated backwards around the x-axis at *M* rotation angles *φ*_*j*_, thus obtaining the rotated truncated tomogram 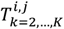. **(c)** For each 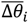 and *φ*_*j*_, the rotated truncated tomogram 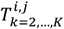 is reprojected along the *z*-axis, thus obtaining the reprojected QPMs 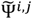, **(d)** which are compared to the experimental QPM Ψ_1_ at *θ*_1_ = 0° by the TSI **(e)** to build the functional 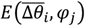, which minimum is highlighted in green. **(f)** After computing the filtered functional 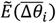, the average angular increment 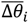 is found by minimizing it (green dot), **(g)** which is finally used along with the positions *y*_*k*_ and the average translational increment 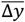 to retrieve the unknown rolling angles *θ*_*k*_.

4.1 A search range for the 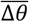 value is fixed, made of *N* values 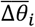 ([0.25°, 12°]in this example with step 0.25°).
4.2 For each 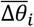, with *i* = 1, …, *N*,
  a. all the corresponding angles 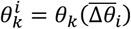 are computed according to Eq. (2);
  b. the truncated tomogram 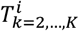 is reconstructed (see Fig. 2(a));
  c. the truncated tomogram 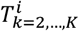 is rotated backwards by an angle *φ*_*j*_ taken from a fixed reprojection range, made of *M* angles *φ*_*j*_, thus obtaining the rotated truncated tomogram 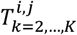 ([0.25°, 12°] in this example with step 0.25°, see Fig. 2(b));
  d. for each reprojection angle *φ*_*j*_, with *j* = 1, …, *M*,
    i. the rotated truncated tomogram 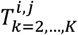 is reprojected along the optical *z*-axis to emulate the QPM formation, thus obtaining the reprojected QPM 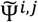 (see Fig. 2(c));
    ii. the reprojected QPM 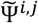 is compared to the first experimental QPM Ψ_1_ by the TSI metric (see Fig. 2(d)).
4.3 A functional 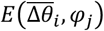 is built, with *i* = 1, …, *N* and *j* = 1, …, *M* (see Fig. 2(e)).
4.4 The functional 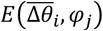 is averaged with respect to the *φ* variable for each *i* = 1, …, *N*, i.e.

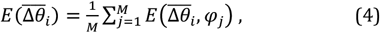
4.5 A moving average with length 3 is applied to the functional 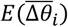, thus obtaining the filtered functional 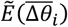 (see Fig. 2(f));
4.6 The 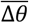 value is finally computed by minimizing the filtered functional 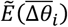 (see Fig.2(f)). The filtered functional 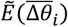 in Fig. 2(f) is minimum in the correct 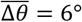, thus the unknown rolling angles can be finally estimated through Eq. (2), as reported in Fig. 2(g). Instead, by using in this example the matching-based method with *f*_2_ = 61 in the Eq. (1), there would be a percentage error of 12.5% respect to the simulated rolling angles 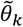, calculated as

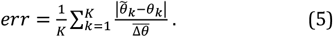

As for HTFC experiments, the opto-fluidic recording system in Fig. 3(a) has been employed, based on a digital holographicmicroscope in off-axis configuration [11,16]. An example of recorded digital hologram containing ovarian cancer cells (CAOV3 cell line) is shown in Fig. 3(b), where the *xyz* Cartesian reference system has been reported. From each digital hologram, ROIs containing the flowing cells have been cropped and converted into the corresponding QPMs (step 1, see Fig. 3(c)) [11,16]. The cell’s transversal positions have been then computed for each frame through holographic tracking algorithms (steps 2-3) [11,16]. To retrieve the unknown rolling angles, the matching-based method can be implemented. However, as shown in Fig. 4(a), a human visual analysis of the QPMs sequence and the TSI trend is requested to detect *f*_2_ = 41 as the frame related to a 360° rotation. Instead, to implement the reprojection-based method (steps 4.1-4.3), we have fixed a first search range [0.25°, 12°] with step 0.25°, thus obtaining the functional 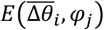 reported in Fig. 4(b) by using all the *K* = 120 experimental QPMs. The 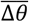 value minimizing the functional 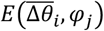 in Fig. 4(b) is 1.75°, which instead should be close to 9° according to the matching-based method in Fig. 4(a). However, the average and filtering operations (steps 4.4-4.6) allow overcoming this error, thus reaching the correct 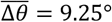 solution by minimizing the filtered functional 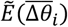 in Fig. 4(c). Furthermore, to refine the 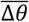 value, the same pipeline can be repeated by considering a narrower and thicker search range and reprojection range around 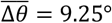, i.e. [8.25°, 10.25°] with step 0.05° (see Fig. 4(d)), and by using only the first *K* = 42 experimental QPMs, that correspond to the frames containing a full 360° rotation according to the Eq. (2) with 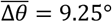. In this way, a refined solution is found, i.e.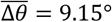. By applying Eq. (2) with 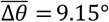, the unknown rolling angles *θ*_*k*_ are retrieved (step 5), as displayed in green in Fig. 4(e). Hence, we have automatically found that a full rotation occurs between frames 40 and 41 (*θ*_40_ = 356.4° and *θ*_41_ = 365.9°), which agrees with the matching-based solution obtained through the human inspection, i.e. *f*_2_ = 41, but is more accurate as the *ε* error is avoided, as reported in red in Fig. 4(e). Finally, using the retrieved rolling angles *θ*_*k*_, the 3D RI tomogram of the CAOV3 cell has been reconstructed (see Fig. 4(f)).

**Fig. 3.**
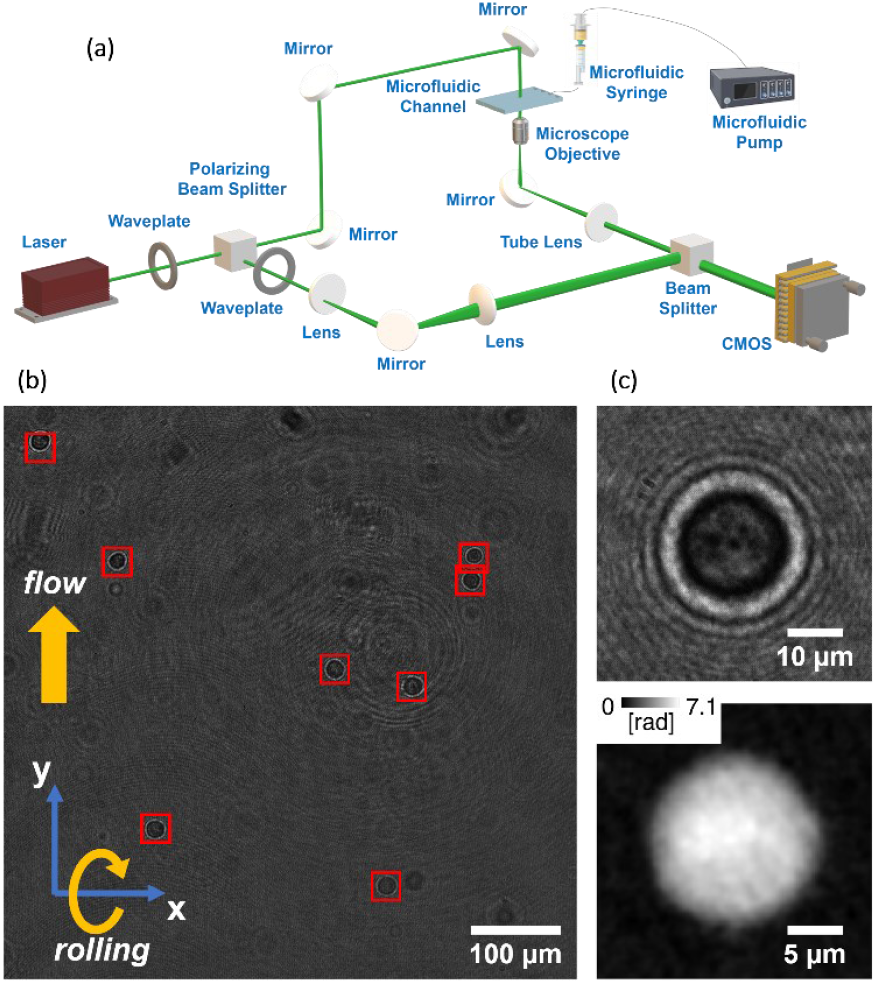
HTFC experiments of CAOV3 cells. **(a)** Opto-fluidic recording system. **(b)** Digital hologram taken from a recorded video sequence, with highlighted in red the ROIs containing the detected and tracked cells. **(c)** At the top, holographic ROI; at the bottom, corresponding QPM.

**Fig. 4.**
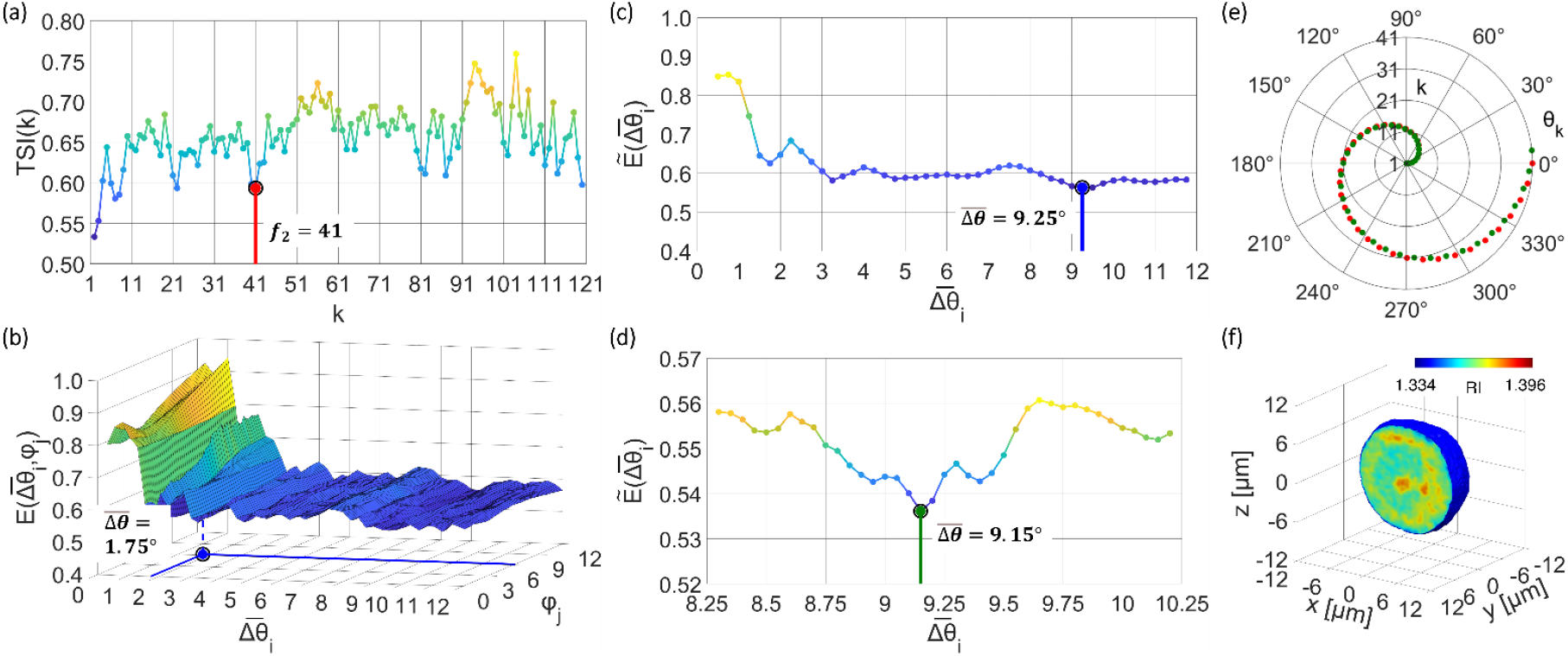
Retrieval of the unknown rolling angles of a CAOV3 cell. **(a)** Matching-based method, with highlighted in red the *f*_2_ frame found by human inspection. **(b-d)** Reprojection-based method. In (b), the functional is minimum in the wrong 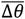 (blue dot); in (c), this error is corrected by filtering the functional (blue dot); in (d), the average angular increment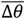 is refined (green dot) after narrowing and thickening the filtered functional in (c). **(e)** Comparison between the unknown rolling angles retrieved through the matching-based method in (a) (red) and the reprojection-based method in (d) (green). **(f)** Central slice of the 3D RI tomogram reconstructed using the angles retrieved by the reprojection-based method in (e).

To implement the reprojection-based method and recover the unknown rolling angles, we have utilized the integral QPM reprojection and the Filtered Back Projection algorithm for tomographic reconstruction [11,16]. Of course, the employment of more advanced QPM reprojection algorithms [17,18] or tomographic reconstruction algorithms [19] is expected to provide an even more accurate result, but at the cost of a greater computational burden. In fact, in the configuration herein proposed, the reprojection-based method has allowed to significantly speed up the rolling angles retrieval with respect to the matching-based method, as human intervention has been completely avoided.

Moreover, it is worth underlining that the computational time and the accuracy of the reprojection-based method can be further enhanced by narrowing the search range of the optimization problem thanks to the availability of prior information about an approximate average angular increment, provided for example by a precise microfluidic control. In conclusion, we introduced a self-consistent reprojection-based method for retrieving the unknown 3D rotations of single cells in HTFC. By iteratively optimizing the alignment between experimental and reprojected QPMs, our approach enables an automatic and accurate RI tomographic reconstruction of flowing cells. This advancement represents a significant step toward 3D label-free imaging of single cells in flow cytometry by allowing highly reliable HTFC and expanding the practical applicability of this promising technology in biomedical research and diagnostics.

## Acknowledgment

This work was supported by project PRIN 2022 PNRR - Flow-cytometry ImaGing by Holographic tomography for predicting TUMor control in Oncology patients treated with Radiotheraphy (FIGHT-TUMOR), Prot. P2022ATE2J – funded by the Italian Ministry of University & Research in the framework of Next Generation EU.

